# The characterisation of antimicrobial resistant *Escherichia coli* from dairy calves

**DOI:** 10.1101/2023.04.03.533045

**Authors:** Merning Mwenifumbo, Adrian L Cookson, Shengguo Zhao, Ahmed Fayaz, Jackie Benschop, Sara A Burgess

## Abstract

Dairy calves, particularly pre-weaned calves have been identified as a common source of multidrug (MDR) resistant *E. coli*. However, the strains and whether their resistance genes are plasmid or chromosomally located have not been well characterised. Our study examined the phenotype and genotype of antimicrobial resistant *E. coli* isolated from young calves (≤ 14 days old). Recto-anal swab enrichments from 40 dairy calves located on four dairy farms were examined for tetracycline, streptomycin, ciprofloxacin, and third-generation cephalosporin resistant *E. coli*. Fifty-eight percent (23/40) of calves harboured antimicrobial resistant *E. coli*: 18/40 (45%) harboured tetracycline resistant and 25% (10/40) harboured chromosomal mediated AmpC producing *E. coli*. Whole genome sequencing of 27 isolates revealed five sequence types, with ST88 being the dominant ST (17/27, 63% of the sequenced isolates) followed by ST1308 (3/27, 11%), along with the extraintestinal pathogenic *E. coli* lineages ST69 (3/27), ST10 (2/27, 7%), and ST58 (1/27, 4%). Additionally, 16 isolates were MDR, harbouring additional resistance genes that were not tested phenotypically. Oxford Nanopore long-read sequencing technologies enabled the location of multiple resistant gene cassettes in IncF plasmids to be determined. A phylogenetic comparison of the ST10 and ST69 isolates demonstrated that the calf derived isolates were distinct from other New Zealand animal, human, and environmental isolates. and highlights the importance of understanding the sources of antimicrobial resistance.

## 1. Introduction

Antimicrobial resistance (AMR) has become a global human, animal and environmental health concern threatening current effective prevention and treatment options. The spread of antimicrobial resistant bacteria and their genes is complex and multifaceted, with one of the main drivers being the use of antimicrobials in humans and animals. Although current evidence indicates that human reservoirs are the main sources for the spread of resistant bacteria within human populations, the emergence and transmission of resistant strains does occur via other pathways including contact with animals and ingestion of food products.

The use of antimicrobials in livestock has come under critical review, due to the association of “routine” antimicrobial use in livestock with the emergence and spread of AMR. Antibiotic use in food-producing animals is low in New Zealand compared to the United States, Canada, Australia, and many European countries (Hillerton et al., 2016). Generally, low antibiotic use is reported in cattle, compared to other food animals such as swine or poultry (Marshall Bonnie and Levy Stuart, 2011). In the dairy sector, antibiotics are used to treat a variety of infections including endometritis, mastitis, foot-rot, and respiratory infections (Burgess and French, 2017). Of particular concern are the 3^rd^ and 4^th^ generation cephalosporins which have been associated with a dairy farm being positive for ESBL and AmpC producing *Enterobacteriaceae* (Gonggrijp et al., 2016; Hordijk et al., 2019; Snow et al., 2012). The use of other antimicrobials may also be important for the selection of other AMR determinants, including multidrug resistant *E. coli*. Other farm management practices, such as waste milk feeding and housing, may also be associated with the emergence and spread of antimicrobial resistant bacteria (Collis et al., 2019; Springer et al., 2018).

Young calves (up to two weeks in age) are more likely to shed antimicrobial resistant *E. coli*, compared with older calves or lactating cows (Awosile et al., 2018; Berge et al., 2010; Hordijk et al., 2013a; Weber et al., 2021). Additionally, the abundance of resistance genes decreases in the gut of calves over time (Liu et al., 2019). Feeding waste milk to calves has been associated with changes in the calf microbiome, and an increase in antimicrobial resistant bacteria and their genes (Maynou et al., 2017b; Penati et al., 2021; Pereira et al., 2014). Waste milk is milk from cows that have been treated with antibiotics or other types of drugs before the withholding period is over. Waste milk also refers to milk with increased somatic cell counts (over 150,000 cells/ml) (Ricci et al., 2017). Studies have found that waste milk containing fourth generation or third generation cephalosporins, was associated with the presence of third generation cephalosporin resistant *E. coli* (Aust et al., 2013; Brunton et al., 2014; Randall et al., 2014). Additionally, other types of antimicrobial resistant *E. coli* have been associated with waste milk feeding such as chloramphenicol, aminoglycoside, and beta-lactam resistance (Duse et al., 2015; Maynou et al., 2017b). However, the genetic basis for these resistance types of *E. coli* has not been well described. Here we used whole genome sequencing to characterise multidrug resistant *E. coli* isolated from calves located on dairy farms that fed waste-milk to their replacement calves and explored the genetic relationship of these *E. coli* with New Zealand isolates and other calf and dairy cattle isolates.

## 2. Methods

### 2.1 Isolation of E. coli and antibiotic susceptibility testing

*E. coli* was isolated from previously collected recto-anal mucosal swabs from calves (2-14 days old) enriched in modified Tryptone Soy Broth (Browne et al., 2018), by streaking onto four agar plates: MacConkey agar, two selective MacConkey agar plates (each containing 1 mg L^-1^ cefotaxime sodium (Sigma Aldrich, St. Louis, MO, USA) or 1 mg L^-1^ ceftazidime pentahydrate (Sigma Aldrich) as well as CHROMagar™ ESBL (CHROMagar, Paris, France) and incubated at 35°C overnight. Presumptive *E. coli* isolates were purified and identified as previously described (Burgess et al., 2021a). Confirmed *E. coli* isolates were screened against the six antibiotics cefoxitin (30 µg), cefpodoxime (10 µg), cefotaxime (5 µg), tetracycline (30 µg), streptomycin (10 µg) and ciprofloxacin (5 µg) using the Kirby-Bauer disc diffusion tests with both EUCAST and CLSI guidelines (Table S1). An AmpC confirmation disc diffusion test was carried out on those *E. coli* that were resistant to cefoxitin and an ESBL confirmation disc diffusion test was carried out on those *E. coli* that were resistant to cefpodoxime and/or cefotaxime as previously described (Toombs-Ruane et al., 2020).

### 2.2 Molecular characterisation

Crude DNA extractions were prepared using the heat lysis method where 3-4 colonies were suspended in 1ml of nuclease-free water in an Eppendorf, heated at 100°C for ten minutes, and the supernatant was stored at −20°C for later PCR reactions. Confirmed antimicrobial resistant *E. coli* isolates were phylotyped by quadraplex PCR as described by (Clermont et al., 2013). Confirmed AmpC producing isolates were further analysed by PCR to determine if they harboured genes encoding the plasmid-mediated AmpC enzymes (types MOX, CIT, DHA, ACC, and FOX) as described by (Pérez-Pérez and Hanson, 2002), or if they harboured a mutation in the promoter region of the *ampC* chromosomal gene as described by (Caroff et al., 1999).

### 2.3 DNA extraction, whole genome sequencing and assembly

*E. coli* isolates that were resistant to two antibiotics (tetracycline, streptomycin, or cefoxitin) were selected for short-read whole genome sequencing. Six of these isolates were also sequenced using long-read Oxford Nanopore Technologies (ONT). For short-read sequencing genomic DNA was extracted using the QIAamp DNA Mini Kit (Qiagen, Hilden, Germany) as per the manufacturers’ instructions. The DNA was eluted in 50µl of sterile PCR-grade water. The library preparations and next-generation sequencing were performed by either Massey Genome Sequence (Massey University, Palmerston North, New Zealand) using the Illumina MiSeq™ (San Diego, California, U.S.A) or Beijing Genomics Institute (BGI, Qingdao, China) using the BGI DNBSEQ™. The Illumina Nextera™ XT library preparation kit (San Diego, California, U.S.A) was used to prepare the libraries as per the manufacturers’ instructions and sequencing was performed on an Illumina MiSeq™ (San Diego, California, U.S.A). For sequencing using the BGI DNBSEQ platform, DNA was sheared into 250- to 300-bp fragments using a Covaris M220 Focused-Ultrasonicator (ThermoFisher Scientific, Waltham, MA, United States, and libraries were then prepared using the MGIEasy Universal DNA Library Prep Set (MGI, Shenzhen, China).

Raw DNA sequence reads were quality assessed using FastQC (v. 0.11.9) (Andrew, 2010), then trimmed and quality filtered using fastp (v. 0.11.9) (Chen et al., 2018), prior to assembly using SKESA (v. 2.3.0) as part of the Nullarbor pipeline (v. 2.0.20191013) (Seemann et al., 2016; Souvorov et al., 2018), using *E. coli* JJ1887 (accession CP014316) as the reference (Johnson et al., 2016).

For long-read sequencing, genomic DNA was extracted using the Wizard® Genomic DNA Purification Kit (Promega, Madison, WI, USA) and the libraries were prepared using the Rapid Barcoding Sequencing kit (SQK-RBK004; ONT, UK) as previously described (Collis et al., 2022). The libraries were loaded onto a SpotON flow cell R9 version FLO-MIN106D and sequenced using a MinION Mk1B device (ONT, Oxford, England). Base-calling was carried out using Guppy (v. 6.1.2) with the super high accuracy model. Reads were demultiplexed using qcat (v. 1.1.0; https://github.com/nanoporetech/qcat), trimmed using Porechop (v.; https://github.com/rrwick/Porechop) and filtered using Filtlong (v. 0.2.1; https://github.com/rrwick/Filtlong), keeping a minimum length of 1,000bp. The reads were then assembled using either Flye (v. 2.9) (Kolmogorov et al., 2019) or Unicycler (v. 0.4.4) (Wick et al., 2017). Flye assemblies were polished with short-reads, using three rounds of Pilon (v. 1.24) (Walker et al., 2014). All genome data for this study have been deposited in GenBank under the Bioproject accession PRJNA938096.

For all the genome assemblies the resistance genes, plasmid type and serotype were determined using ABRicate (v. 1.0.0) (Seemann, 2020) with the databases (all updated 2022-Sep-11).: ResFinder (Zankari et al., 2012), CARD (Alcock et al., 2019), PlasmidFinder (Carattoli et al., 2014), and EcOH (Ingle et al., 2016).

### 2.4 Genome sequences and bioinformatics

The sequence reads of other New Zealand ST10, ST69 strains as well as ST88 cattle and calf isolates previously sequenced by other institutes (Table S1) were downloaded from the NCBI Sequence Read Archive database and processed using the Nullarbor pipeline (v. 2.0.20191013). Whole genome multi-locus sequence typing (wgMLST) was undertaken using Fast-GeP (v. 1.0) (Zhang et al., 2018) and Single Nucleotide Polymorphism (SNP) analyses using Snippy (v. 4.3.6) (Seemann, 2015). Neighbour-joining trees were generated using the resulting Snippy alignments in SplitsTree (v. 4.1.7.1) (Bryant and Moulton, 2004) and the final trees were visualised using the Interactive Tree of Life (iTOL v. 6.7) (Letunic and Bork, 2021).

### 2.5 Plasmid analysis

The closest plasmid relatives were determined using the plasmid database, PLSDB (v. 2021_06_32), using the default settings (Galata et al., 2018). Plasmid and resistance cassette comparisons were carried out in EasyFig (v.2.2.2) (Sullivan et al., 2011). Plasmid incompatibility and sequence types were determined using pMLST (v. 2.0) and PlasmidFinder (v. 2.0) (Carattoli et al., 2014).

## 3. Results

Forty recto-anal mucosal swab-enrichments, previously collected from 2 to 9-day old calves (dairy replacement, bobby, and beef calves; Table S2) located on four farms (Table 1), were screened for ESBL- and AmpC-producing, as well as tetracycline-, streptomycin- and ciprofloxacin-resistant *E. coli.* No *E. coli* were isolated from one calf from farm VCF77. Overall, 58% (23/40) of the calves harboured antibiotic resistant *E. coli.* Of these 23 calves, 18 harboured tetracycline resistant *E. coli*, 13 harboured streptomycin resistant *E. coli* and 10 harboured AmpC-producing *E. coli* (Figure 1). Additionally, 16 calves harboured *E. coli* that were co-resistant, to tetracycline, streptomycin and/or AmpC-producing (Figure 1). None of the calves harboured ESBL-producing or ciprofloxacin resistant *E. coli*.

**Table 1.** Description of farms

| Farm sample ID | Date sampled | Milking herd size | Breed | Other livestock | Replacement calves isolated | Colostrum |
| --- | --- | --- | --- | --- | --- | --- |
| VCF77 | 2014-09-11 | 950 | Friesian | No | Yes | Yes |
| VCF79 | 2014-09-15 | 1100 | Friesian/Jersey or Jersey | Pigs and sheep | No | Yes |
| VCF80 | 2014-09-15 | 1000 | Friesian | Pigs | Yes | Yes |
| VCF89 | 2014-09-17 | 1700 | Friesian or Holstein | Pigs and sheep | Yes | Yes |

**Figure 1.**
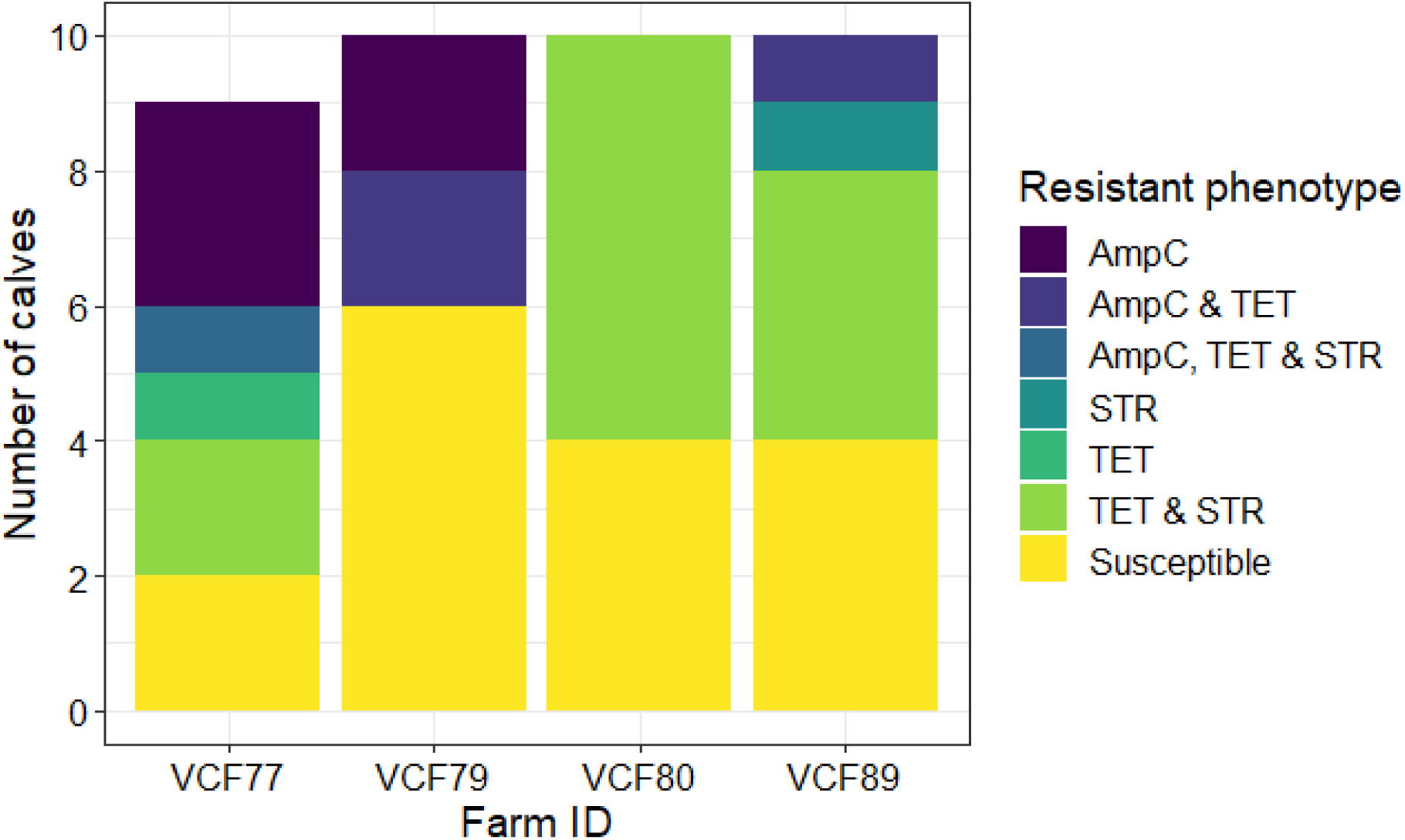
Number of calves harbouring isolates that showed various resistance phenotypes from the four farms.

### 3.1 Genotypic characterisation of the antimicrobial resistant *E. coli*

A total of 57 resistant *E. coli* isolates from 23 calves were phylotyped using the Clermont method (Clermont et al., 2013). Results showed that the resistant *E. coli* isolates were distributed among phylogroups B1, C and D (Table 2). All the AmpC-producing *E. coli* were confirmed to have mutations in the promoter and attenuator regions of the chromosomal *ampC* gene at positions at positions −42, −18, −1 and +58, and originated from farms VCF77 or VCF89.

**Table 2.** Distribution of E. coli phylogroups among resistant E. coli isolates

|  | Number of isolates by phylogroup (No. of calves) |  |  |  |  |  |
| --- | --- | --- | --- | --- | --- | --- |
|  | AmpC only | AmpC and TET | TET and/or STR resistant |  |  | Total |
|  | B1 | C | B1 | C | D |  |
| VCF77 | 12 (4) |  |  | 2 (2) | 3 (3) | 17 (9) |
| VCF79 |  |  | 1 (1) | 6 (5) |  | 7 (6) |
| VCF80 |  |  | 2 (2) | 8 (6) | 1 (1) | 11 (9) |
| VCF89 |  | 8 (2) | 2 (2) | 7 (5) |  | 17 (9) |
| **Total** | **12 (4)** | **8 (2)** | **5 (5)** | **23 (18)** | **4 (4)** |  |

### 3.2 Population structure of AmpC-producing, tetracycline resistant and streptomycin resistant *E. coli*

Twenty-seven *E. coli* isolates displaying various AMR phenotypes were selected for whole genome sequencing to explore their genetic relatedness and diversity (Table 3, Figure 2). The whole genome sequence data were used to determine sequence type (ST), serotype, plasmid type, and AMR genes. Most of the isolates belonged to ST88 (17/27, 63%) with the remaining belonging to ST69 (3/27, 11%), ST1308 (3/27, 11%), ST10 (2/27, 7%), and ST58 (2/27, 7%). There was variability in the ST of strains isolated from each farm. Farm VCF77 had the least variability with only two STs isolated.

**Table 3.**
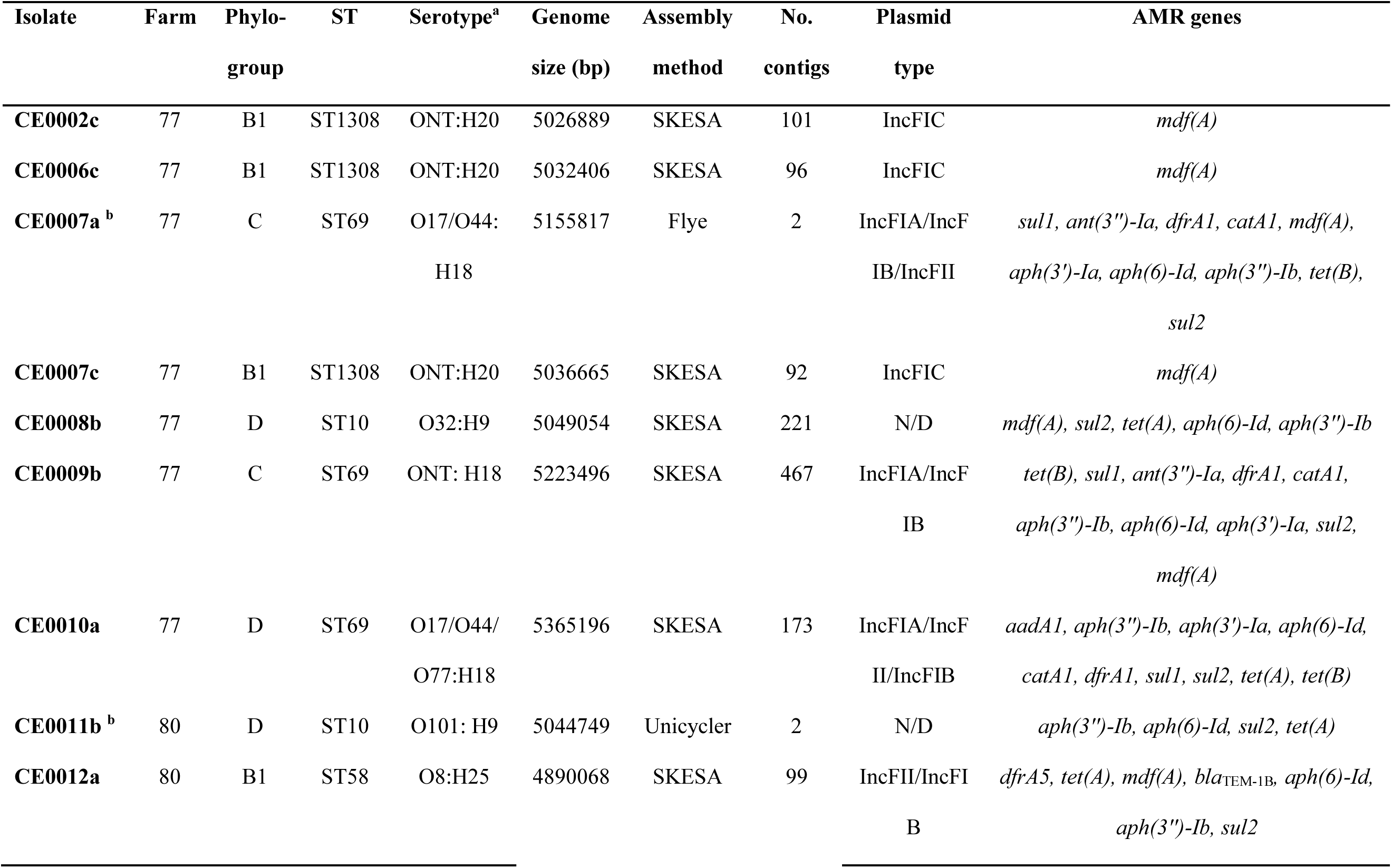
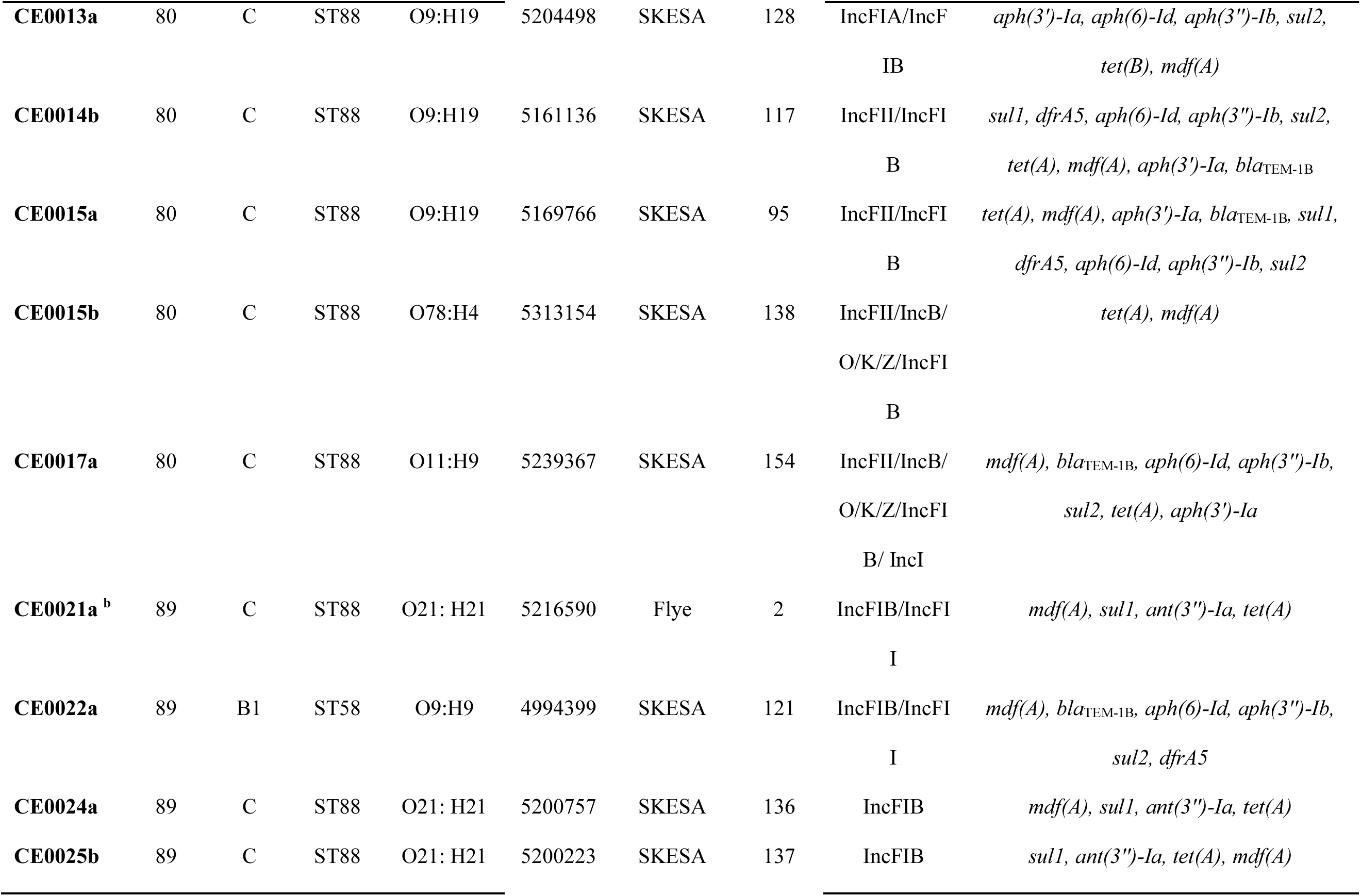
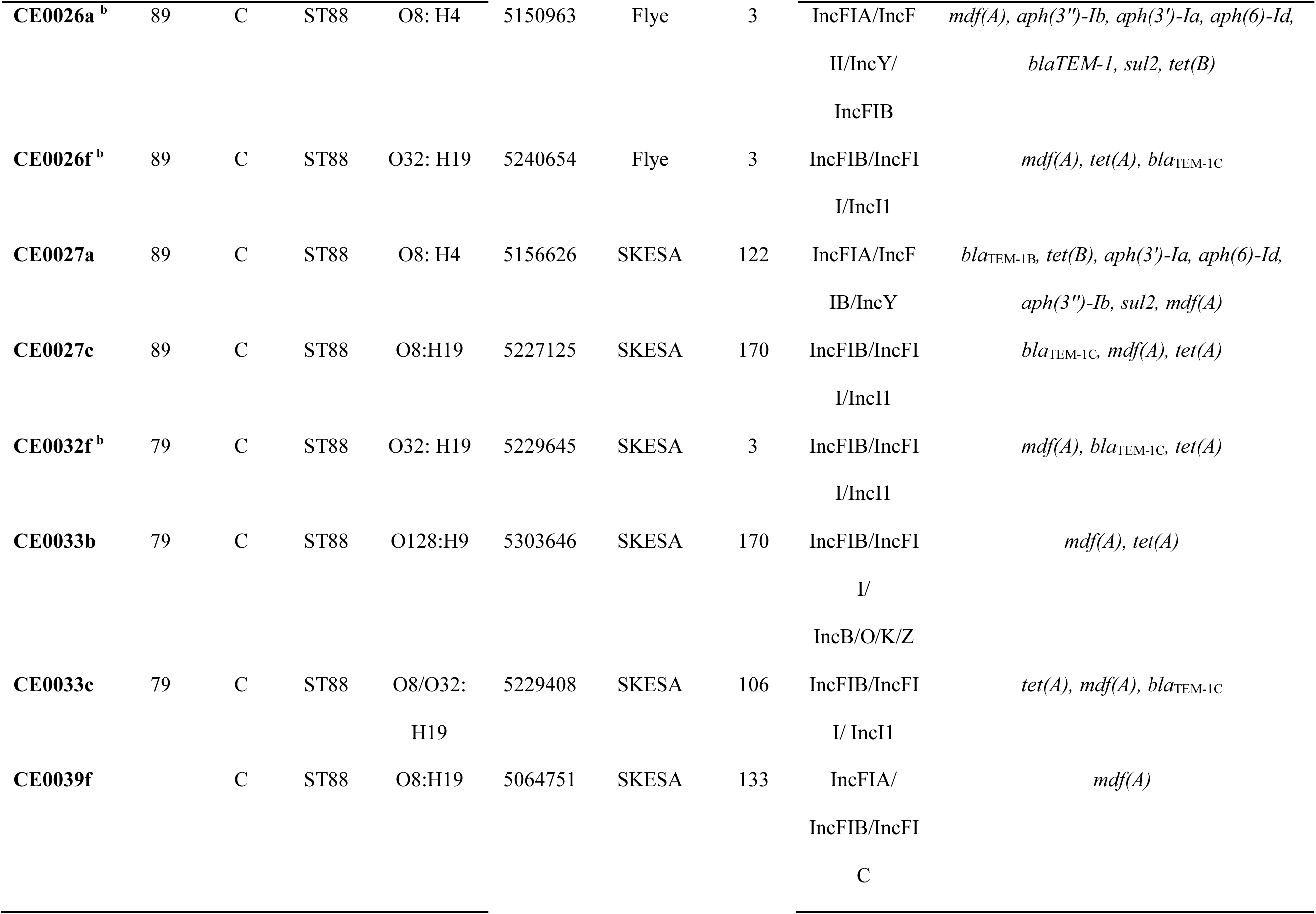
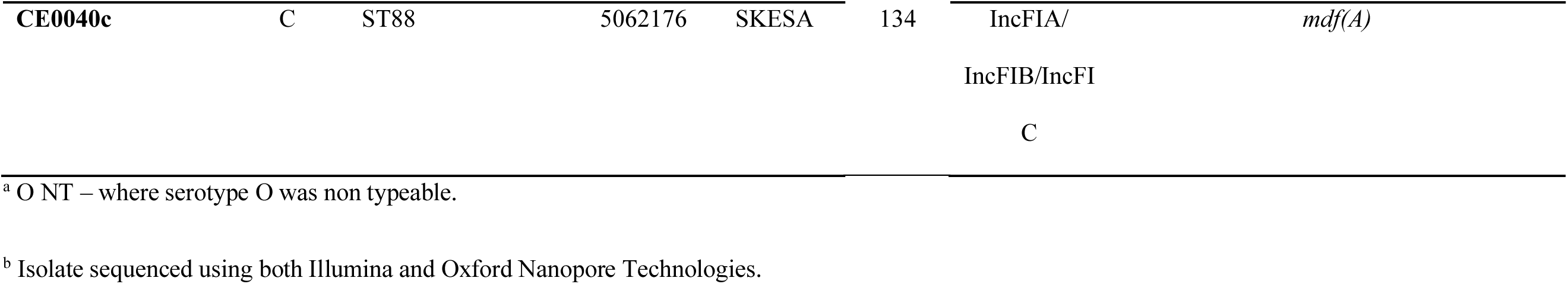
Genomic characteristic of the 27 *E. coli* isolates

**Figure 2.**
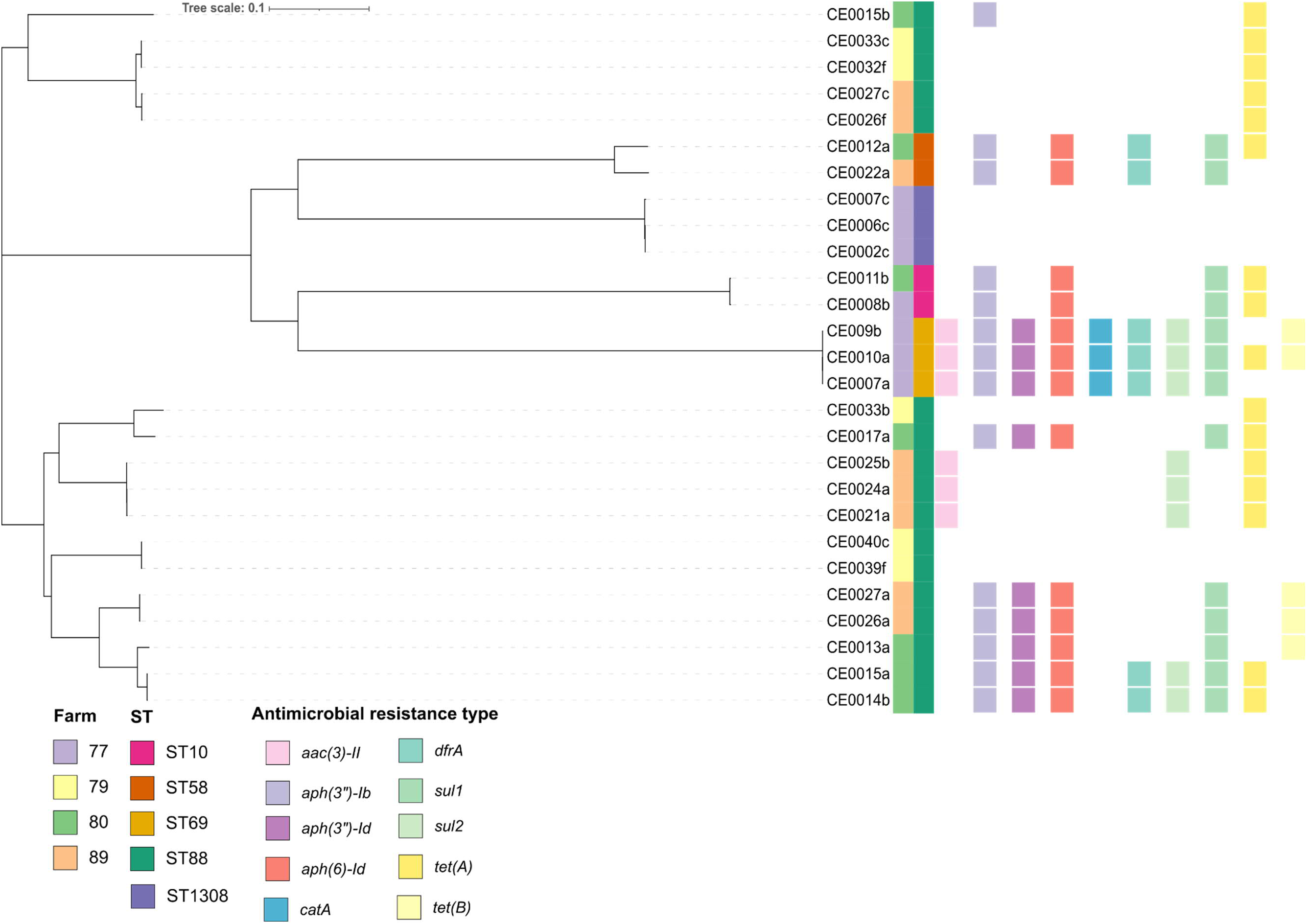
Neighbour joining tree of 27 *E. coli* isolated from New Zealand dairy calves, generated from 3,294 shared alleles identified using the wgMLST tool Fast-Gep. The first coloured strip represents the farm, the second strip the sequence type, followed by the presence (coloured squares) or absence of antimicrobial resistance genes.

The AMR genotype was concordant with the phenotypic results. Noticeably no plasmid mediated AmpC beta lactamase and ESBL genes were found. However, some additional AMR genes, *catA1* for chloramphenicol resistance, *dfrA1* and *dfrA5* for trimethoprim resistance, *sul1* and *sul2* for resistance to sulphonamides and *qacEdelta1* for resistance to quaternary ammonium were detected. These were not tested for phenotypically. Based on their genotype, 16 of 27 isolates were MDR (defined as having ARGs conferring resistance to three or more classes of antibiotics). The ST69 isolates harboured the greatest number of resistance genes (8-10 ARGS). No ARGs were detected in the ST1308 genomes. In general, each clonal group harboured the same resistance determinants. Exceptions were the ST69 isolates CE0007a, CE009a and CE0010a which had differences in the number and type of genes conferring resistance to tetracycline as well as the ST88 isolate CE0033b where only two resistance genes (*mdf(A)* and *tet(A)*) were detected.

To give insights into the transmission of these resistant *E. coli* from a One Health perspective a core SNP analysis was undertaken on the ST10 and ST69 isolates (Figure 4, Supplementary Figure1, and Table S3) (Burgess et al., 2022). The New Zealand dairy calf isolates from this study were distinct from the other New Zealand derived isolates but were closely related to each other, with the ST10 isolates from farms VCF77 and VCF80 differing by six SNPs suggesting they were derived from a shared source or that there was farm to farm transmission (Table S3).

**Figure 3.**
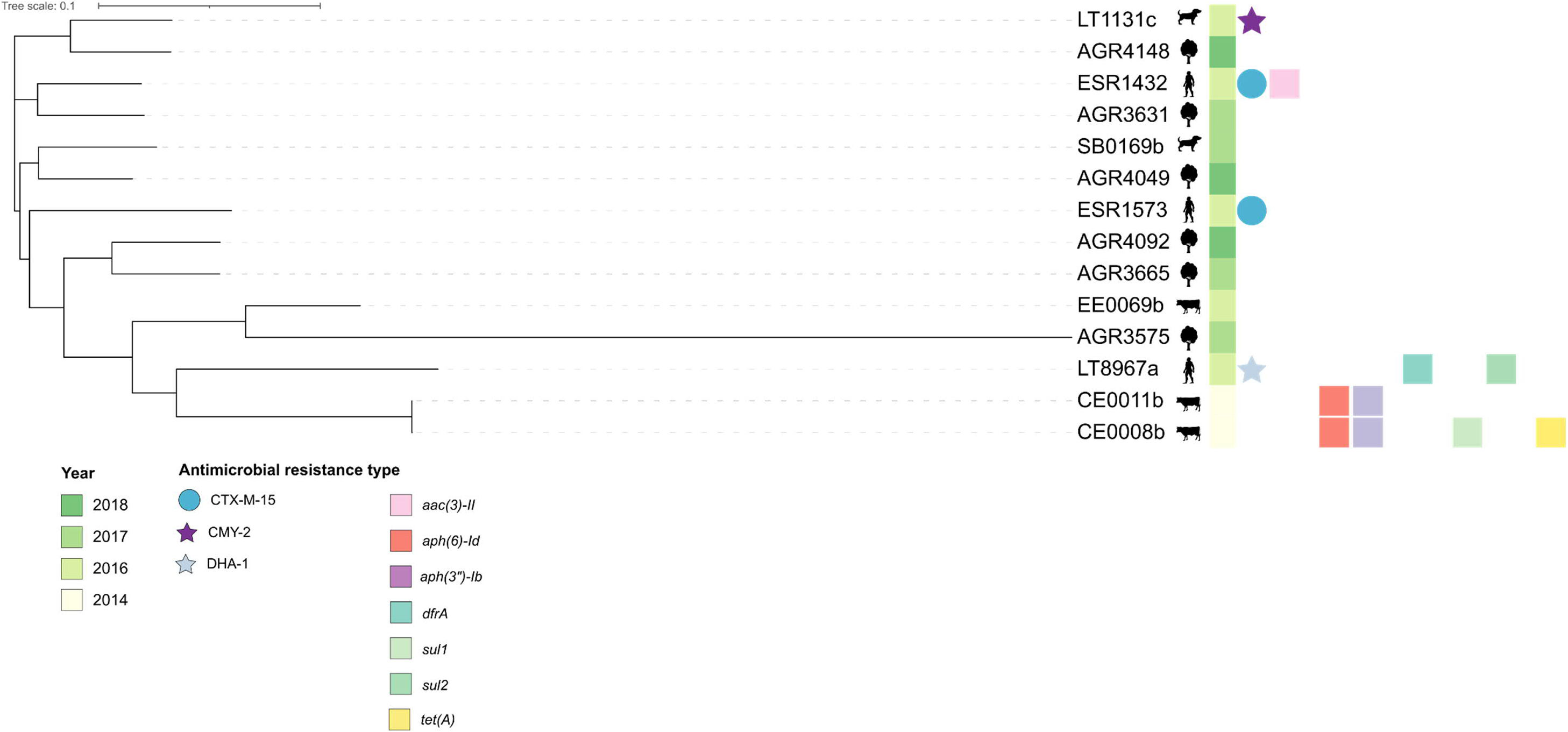
Neighbour joining phylogeny of New Zealand *E. coli* ST10 isolates generated using 30,198 core SNPs. CE0011b was used as the reference. The symbols represent the source (dog, natural environment, human, or cattle/calf) of the isolate, the coloured strip represents the year of isolation, the circles represent the presence of the *bla*CTX-M gene, the stars the presence of the *bla*CMY or *bla*DHA-1 gene and the squares the presence of other antimicrobial resistance genes.

**Figure 4.**
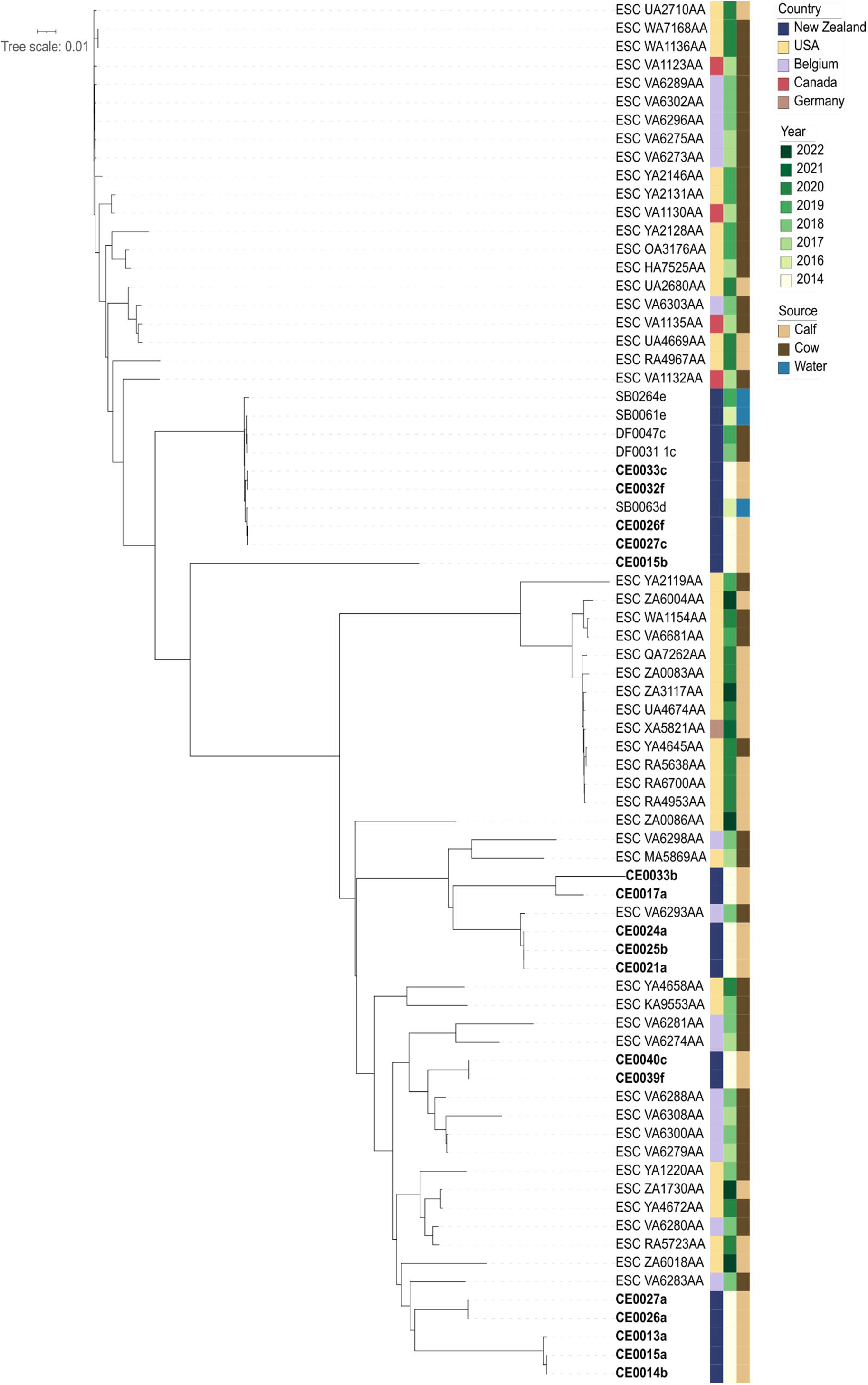
Neighbour joining phylogeny of *E. coli* ST88 isolated from calves and adult cattle, generated using 19, 422 core SNPs. CE0026f was used as the reference. The first coloured strip represents the country of isolation, the second strip represents the year of isolation, and third strip represents the source.

*E. coli* ST88 from New Zealand have rarely undergone whole genome sequence analysis, but this sequence type has frequently been isolated from cattle, therefore, we compared the 17 ST88 isolates with other ST88 adult cattle and calf isolates (Figure 5). Our New Zealand calf derived isolates were distributed throughout the phylogeny but generally clustered by farm. There was one exception where isolates from farms VCF89 and VCF79 (CE0026f, CE0027c, CE0032, and CE0033) differed by 3-46 SNPs.

**Figure 5.**
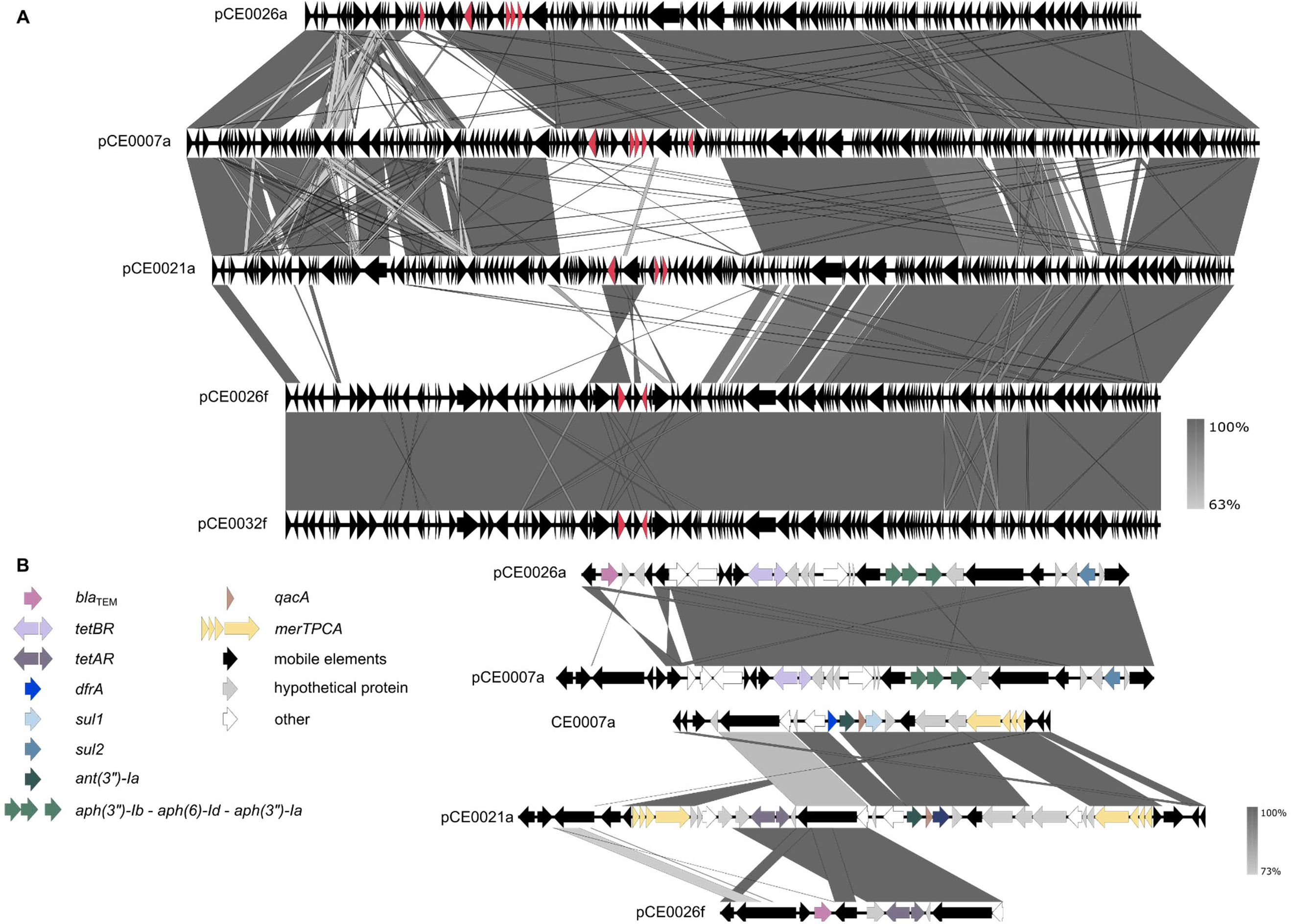
Comparison of IncF plasmids and their resistance gene cassettes. (A) Genetic maps and degree of homology between the IncF plasmids from five strains of *E. coli* isolated from New Zealand dairy calves. (B) Genetic maps and degree of homology between the resistance gene cassettes.

None of the isolates could be classified as STEC and 23 were classified as ExPEC, containing at least two of the genes *afaA*, *iutA*, *kpsM*, and *papC* (Joensen et al., 2014; Johnson et al., 2018; Malberg Tetzschner Anna et al., 2020; Sarowska et al., 2019) (Table S4). Eleven isolates contained the gene *astA*, which is encodes the enteroaggregative *E. coli* enterotoxin EAST1 and 23 isolates had at least one gene associated with siderophores (*fyuA, iutA*, and/or *iucC*).

### 3.3. Plasmid analysis

Contigs harbouring plasmid replicons were identified from 25 isolates, of which five were fully assembled using long-read ONT sequencing. IncF was the most common plasmid replicon identified in the 25 isolates, followed by IncI (5/27 isolates), IncB/O/K/Z (3/27 isolates) and IncY (2/27 isolates). The IncF plasmids only harboured resistance genes.

Six isolates, originating from the four different farms, were selected for long-read sequencing to enable the genetic location of the AMR genes to be determined and the structure of the plasmids to be compared (Figure 5). A similar backbone (including the genes required for conjugation) was shared across the five plasmids with differences in the resistance gene cassettes, mobilome and defence mechanisms. Plasmids pCE0026f and pCE0032f were near identical. Three different resistance gene cassettes were identified (Figure 5b). The closest relatives of the five IncF complete plasmids from the long-read assemblies, originated from *E. coli*, *Salmonella enterica* and *Klebsiella pneumoniae* (Table S5) (AbuOun et al., 2021; Zhao et al., 2020).

## 4. Discussion

The main aim of this study was to phenotypically and genotypically characterise antimicrobial resistant *E. coli* isolated from dairy calves located on farms where waste milk feeding is practiced. *E. coli* displaying resistance to two or more classes of antibiotics were isolated from 16/40 calves. In our study phenotypic resistance was characterised for only four classes of antibiotics (beta-lactams, tetracyclines, aminoglycosides and fluoroquinolones). For isolates selected from MacConkey agar with no antibiotics, tetracycline resistance was the most common resistance phenotype displayed. Tetracycline resistant *E. coli* have been frequently isolated from young calves (Astorga et al., 2019; de Verdier et al., 2012), and to our knowledge no farm management practices, including antimicrobial use, have been shown to be associated with the prevalence of these resistant bacteria (de Verdier et al., 2012; Khachatryan Artashes et al., 2004).

Tetracycline resistance genes have frequently been detected in both the gut microbiome of dairy calves and *E. coli* isolated from dairy calves and their environment (Haley et al., 2020; Kyselková et al., 2015; Srinivasan et al., 2008). In our study, tetracycline resistance genes (*tetA* and *tetB*) were the most prevalent. On New Zealand dairy farms tetracyclines are the fourth most used antibiotic class (after penicillins, macrolides and cephalosporins), whereas aminoglycosides (including streptomycin, oleandomycin and neomycin) are rarely used (Bryan and Hea, 2017; Burgess et al., 2021b).

Whole genome sequencing enabled the detection of multiple resistant determinants covering six classes of antibiotics, including resistance genes for trimethoprim, sulphonamides and chloramphenicol were found that were not tested for phenotypically. Although our study identified a variety of different plasmid replicons, only the IncF plasmid type was associated with antimicrobial resistant genes. This contrasts with other studies which have identified a range of plasmid types associated with AMR in *E. coli* isolates from livestock (AbuOun et al., 2021). Given trimethoprim/sulphonamides are rarely used on NZ dairy farms and chloramphenicols are prohibited for food-producing animal use, we hypothesise that the resistance genes for these antibiotics (*dfrA*, *sul1*, *sul2*, and *catA*) were co-selected with other resistance genes. As previously found in other studies these genes were co-located on the same plasmid as aminoglycoside and tetracycline resistance genes as well as the *qacEdelta1* gene, which encodes resistance to quaternary ammonium compounds. Disinfectants are used on dairy farms, including quaternary ammonium compounds which are sometimes used in the acid wash for cleaning milking equipment (Gleeson et al., 2013).

One *E. coli* strain in our study also harboured the *mer* operon on the same plasmid as the *tetA*, *sul1*, and *ant(3”)-Ia* resistance genes. The co-location of tetracycline, sulphonamide and aminoglycoside encoding resistance genes with the mercury operon on IncF plasmids has been previously described (Gaeta et al., 2022; Souza et al., 2022; Zhao et al., 2020) and the presence of the *mer* operon was positively correlated with sulphonamide and aminoglycoside resistance genes, but not tetracycline resistance genes (Souza et al., 2022). Both heavy metals and disinfectants have previously been shown to co-select for resistance genes (Baker-Austin et al., 2006; Gaeta et al., 2020; Mazhar et al., 2021).

Other farm management practices, such as waste milk feeding, may also be associated with the presence of antimicrobial resistant *E. coli* (Carattoli et al., 2014; Duse et al., 2015; Maynou et al., 2017a; Maynou et al., 2017b). In the study by Maynou et al. (2017b) feeding waste milk was associated with beta-lactam and florfenicol resistance, but not tetracycline or aminoglycoside resistance. Waste milk has also been identified as a risk factor for ESBL/AmpC-producing *E. coli* positive calves (Weber et al., 2021). Studies have shown that by adding antimicrobials (penicillin) to waste milk an increased antibiotic concentration is associated with an increased number of antimicrobial resistant faecal bacteria (Berge et al., 2006; Langford et al., 2003; Pereira et al., 2014). Our study investigated calves from farms who fed waste milk only, therefore associations between the use of waste milk and the presence of antimicrobial resistant bacteria could not be made. As with our study many other studies have hypothesised that the presence of antimicrobial resistant bacteria is due to waste milk feeding. However, there are limited studies (due to the lack of controls) that have determined a relationship between waste milk feeding and the presence of specific resistant bacteria or their determinants (Kyselková et al., 2015; Randall et al., 2014). Although quantitative data was and is still lacking on the impact of feeding waste milk to calves, the European Food Safety Authority carried out a qualitative risk assessment and found that there was an increased risk that the feeding of waste milk to calves “will lead to increased faecal shedding of antimicrobial-resistant bacteria” (Ricci et al., 2017). The genomic epidemiology of antimicrobial resistant *E. coli* in waste milk and their potential transfer to calves has not been well studied. Although, ESBL-producing *E. coli* have previously been isolated from waste milk (Randall et al., 2014) and a recent study found both the abundance and diversity of AMR determinants was greater in waste milk compared with bulk milk.

Sixteen *E. coli* isolates from three of the four farms were found to carry *E. coli* with a multidrug resistant (MDR) genotype. Data on antibiotic use on our study farms was not available; therefore, we were unable to determine if low antimicrobial use was associated with no MDR *E. coli* being isolated from farm VCF79. The prevalence of MDR *E. coli* from healthy and sick calves has previously been reported to be high, ranging from 9.8-100% (of isolates from non-selective agar) in healthy calves (de Verdier et al., 2012; Salaheen et al., 2019), and 34-68% in diarrhoeic calves (de Verdier et al., 2012; Orden et al., 2000) from Sweden and the USA. Previous studies have found that the prevalence of antimicrobial resistant *E. coli* and their genes is greater in calves compared with adult cattle (Agga et al., 2022; Geser et al., 2012; Khachatryan Artashes et al., 2004; Weber et al., 2021). Although our study did not directly compare calves with adult cattle, MDR *E. coli* have rarely been isolated from New Zealand adult cattle (Burgess et al., 2021a; Burgess et al., 2021b; Collis et al., 2019). It has been proposed that MDR (to tetracycline, streptomycin and sulfadiazine) *E. coli* have adapted to the calf environment and are therefore more likely to persist and to be excreted compared with susceptible *E. coli* (Khachatryan Artashes et al., 2004).

No ESBL or plasmid mediated ACBL producing *E. coli* were isolated from any of the calves, supporting findings from other studies that New Zealand dairy farms have a low prevalence of these resistant bacteria (Burgess et al., 2021a; Burgess et al., 2021b; Collis et al., 2022). The low prevalence of ESBL/ plasmid mediated ACBL producing *E. coli* could be due to New Zealand dairy farms being low users of antibiotics, particularly 3rd and 4th generation cephalosporins (Bryan and Hea, 2017; Burgess et al., 2021b; Compton and McDougall, 2014). Globally the prevalence of ESBL positive samples from calves range from 13-30% in the USA to 30% in Europe (Agga et al., 2022; Haenni et al., 2014b; Salaheen et al., 2019). In our study those ACBL producing *E. coli* isolated were putative chromosomal mediated, with the mutations found in the promoter region of the *ampC* gene previously described in *E. coli* isolates from both calves and adult cattle (Alzayn et al., 2020; Hordijk et al., 2013b; Santiago, 2019). This present study also found that none of the putative AmpC hyperproducing isolates were multidrug resistant (not resistant to more than three classes of antibiotics). This is in line with previous studies that have showed ESBLs and plasmid mediated AmpC, but not overexpressed chromosomal mediated AmpC, being associated with multidrug resistance (Fernández-Martínez et al., 2021; Toombs-Ruane et al., 2023).

In this study, using the 7-allele house-keeping genes MLST scheme five STs were observed with ST88 being the most common. Various studies have reported ST88 being common in both calves and adult cattle, additionally ST88 has been associated with the enterotoxigenic *E. coli* (ETEC) pathotype and AMR (Alzayn et al., 2020; Awosile et al., 2020; Haenni et al., 2014a). The other STs: ST10, ST69, ST58 and ST1308 have previously been isolated from both humans and livestock (Borges et al., 2019; Day et al., 2019; McKinnon et al., 2018; Seo et al., 2023; Shawa et al., 2021). ST10, ST58, and ST69 are commonly associated with extraintestinal infections in humans (Manges et al., 2019; Riley, 2014). Our study demonstrated that those ST10 and ST69 strains that were previously isolated from the environment or animals (dairy cattle and calves, birds, and dogs) in New Zealand were distributed amongst the human isolates, but with no evidence of any sharing of strains between sources.

In conclusion, our study found a high incidence of tetracycline and streptomycin resistant *E. coli* in dairy calves as has been found in other studies, but no ESBL/pACBL-producing *E. coli* in contrast to other European and US studies where these resistant bacteria are commonly isolated from both calves and adult cattle. Whole genome sequencing enabled the genetic location of the tetracycline and aminoglycoside resistance genes to be determined and demonstrated that 16 of 27 (60%) strains sequenced had a MDR genotype. Further investigations are required to determine whether these MDR *E. coli* are associated with the use of waste milk or other farm management practices.

## Declaration of Competing Interest

The authors declare that they have no known competing financial interests or personal relationships that could have appeared to influence the work reported in this paper.

## Acknowledgments

This work was funded by the New Zealand China Food Protection Network. We gratefully acknowledge A. Springer Browne for collecting the faecal samples, the Ministry of Primary Industries, and the Meat Industry Association for funding this collection, and the farmers who allowed the collection of these samples and ongoing use in research. We also thank Niluka Velathanthiri and Lynn Rogers for support in the laboratory. We wish to acknowledge the use of New Zealand eScience Infrastructure (NeSI) high-performance computing facilities, as part of this research.

---

**Figure S1.**
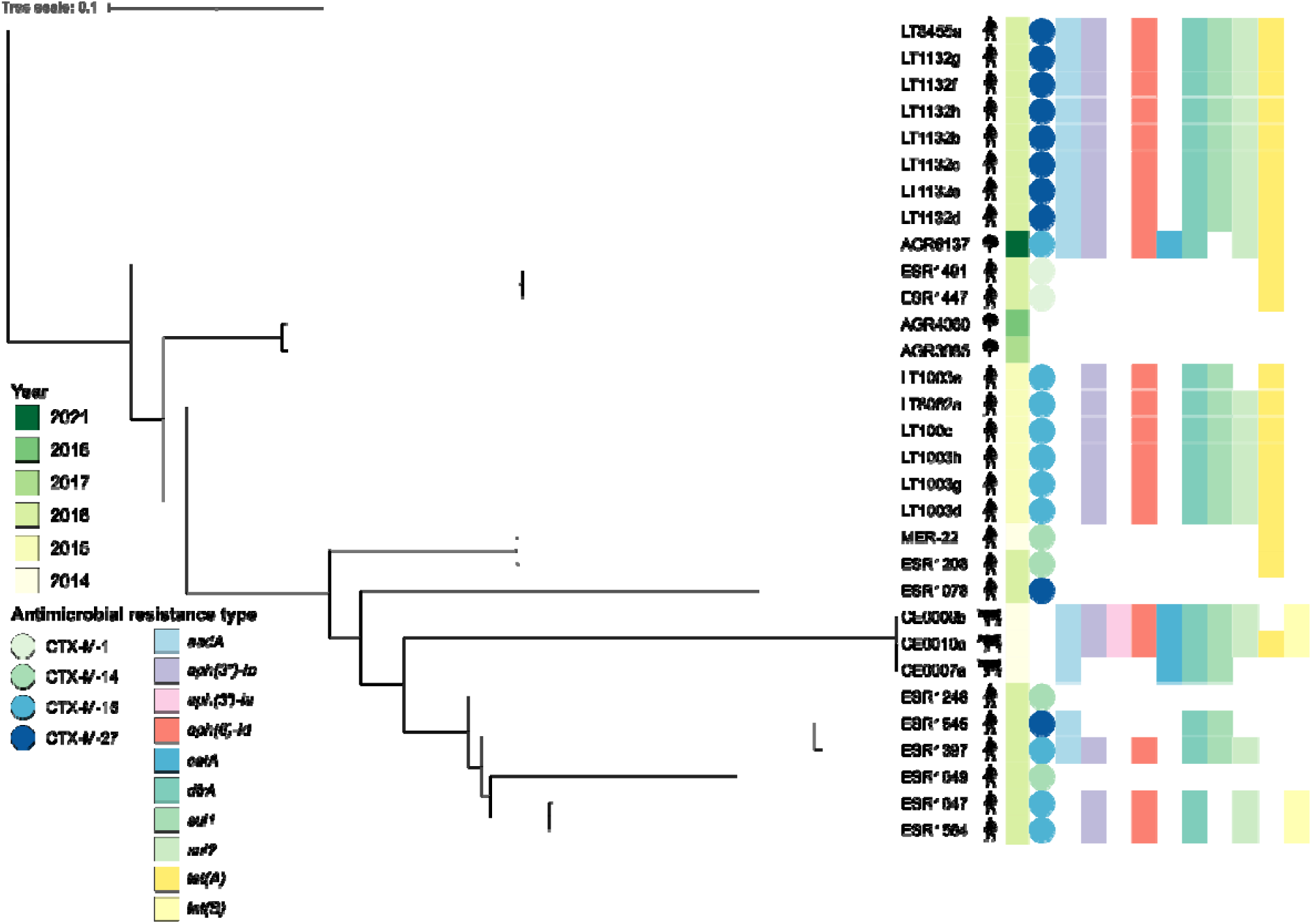
Neighbour joining phylogeny of New Zealand *E. coli* ST69 isolates (adapted from (Burgess et al., 2022) generated using 17218 core SNPs. AGR6137 was used as the reference. The symbols represent the source (dog, natural environment, human, or cattle/calf) of the isolate, the coloured strip represents the year of isolation, the circles represent the presence of the *bla*_CTX-M_ gene, the stars the presence of the *bla*_CMY_ or *bla*_DHA-1_ gene and the squares the presence of other antimicrobial resistance genes. Burgess, S.A., Moinet, M., Brightwell, G., Cookson, A.L., 2022. Whole genome sequence analysis of ESBL-producing Escherichia coli recovered from New Zealand freshwater sites. Microbial Genomics 8.

## Notes

### Competing Interest Statement

The authors have declared no competing interest.

